# Pulmonary Mitochondrial DNA Release and Activation of the cGAS-STING Pathway in Lethal *Stx12* Knockout Mice

**DOI:** 10.1101/2024.10.07.616980

**Authors:** Dan-Hua Liu, Fang Li, Run-Zhou Yang, Zhuanbin Wu, Xiao-Yan Meng, Sen-Miao Li, Wen-Xiu Li, Jia-Kang Li, Dian-Dian Wang, Rui-Yu Wang, Shu-Ang Li, Pei-Pei Liu, Jian-Sheng Kang

## Abstract

STX12 (syntaxin12 or syntaxin13), a member of the SNARE protein family, plays a crucial role in intracellular vesicle transport and membrane fusion. Our previous research has demonstrated that *Stx12* knockout mice exhibit perinatal lethality with iron deficiency anemia. Despite its importance, the comprehensive physiological and pathological mechanism of STX12 remain largely unknown. Here, we uncover that STX12 deficiency causes the depolarization of mitochondrial membrane potential in zebrafish embryos and mouse embryonic fibroblasts. Additionally, the loss of STX12 decreases levels of mitochondrial complex subunits, accompanying mitochondrial DNA (mtDNA) release and activating cGAS-STING pathway and Type I interferon pathway in the lung tissue of *Stx12^−/−^* mice. Additionally, we have observed a substantial increase in cytokines and neutrophil infiltration within the lung tissues of *Stx12* knockout mice, indicating a severe inflammation, which could be a contributing factor for *Stx12^−/−^*mortality. Various interventions have failed to rescue the lethal phenotype, suggesting that systemic effects may contribute to lethality. Further research is warranted to elucidate potential intervention strategies. Overall, our findings uncover the critical role of STX12 in maintaining mitochondrial function and mtDNA stability in pulmonary cells, and reveal that STX12 depletion results in pulmonary mtDNA release and activates mtDNA-dependent innate immunity.

## Introduction

Soluble N-ethylmaleimide-sensitive factor attachment protein receptors (SNAREs) are a family of proteins that play a crucial role in the process of membrane fusion within eukaryotic cells^1,2^. They are essential for various cellular functions, including vesicle trafficking, neurotransmitter release^3,4^, and the fusion of transport vesicles with their target membranes. Syntaxin 12 (STX12) is a member of the Qa-SNARE protein family, which plays a critical role in the endosomal trafficking pathway, where it facilitates the sorting and recycling of endocytosed materials^5,6,7^. The expression of STX12 is reported to be developmentally regulated and is abundant during the late embryonic and early postnatal phases, with levels subsequently diminishing as the brain matures into the late postnatal stages in the mice and rats^8,9^. In addition, in primary neurons and differentiated PC12 cells, STX12 is prominently found in the perinuclear area of the cell body, as well as distributed along neurites and within growth cones^10,9^. The overexpression of STX12 in PC12 cells can enhance the neurite outgrowth during nerve-growth-factor-induced differentiation^11,9^. STX12 is also involved in autophagosome maturation as a genetic modifier of mutant CHMP2B in frontotemporal dementia^12^. Studies have shown that STX12 colocalizes with internalized transferrin and the transferrin receptor^5^. Our previous study has revealed a deficiency in fast recycling of the transferrin receptor in *Stx12^−/−^* MEFs, and demonstrated that *Stx12* knockout mice exhibit iron deficiency anemia and perinatal lethality^13^. However, the pathophysiological alterations and the underlying lethal mechanism of STX12 deficiency remain unknown.

Given the perinatal lethality of *Stx12* knockout mice and the difficulty in exploring potential mechanisms in mice, we generated *Stx12* knockdown zebrafish to investigate the potential effects of STX12 deficiency. Our findings indicated that *Stx12* knockdown in zebrafish resulted in approximately 80% mortality rate within 6 hours post-fertilization, accompanied by mitochondrial damage. Furthermore, we observed a pronounced activation of inflammatory response and reductions in expression levels of mitochondrial complex subunits in the lung tissue of *Stx12* knockout mice. Further mechanistic studies confirmed that mitochondrial DNA (mtDNA) escaped into the cytoplasm, leading to the activation of the cGAS-STING pathway and subsequently triggering the Type I interferon pathway in alveolar epithelial cells of *Stx12* knockout mice. Notably, we discovered that this inflammatory response was closely associated with neutrophils, wherein a positive feedback loop involving neutrophil infiltration and cytokine release exacerbated the inflammation. This amplified inflammatory response is speculated to be a potential cause of perinatal mortality observed in *Stx12* knockout mice.

## Results

### The zebrafish model of *Stx12* knockdown

To gain insights into the function of *Stx12* and the lethal mechanism associated with *Stx12* deficiency, we developed a *Stx12* morpholino (MO) knockdown zebrafish model by embryonic injection of *Stx12*-targeting morpholinos. Two specific morpholino antisense strategies were employed to disrupt the zebrafish *Stx12* gene: one to block translation (ATG-MO) and the other to interfere with the proper splicing of exon 4 (E4I4-MO) (Figure S1A). The effectiveness of *Stx12* knockdown was validated by Sanger sequencing (Figure S1B) and qRT-PCR (Figure 1C). Consistent to our previous findings that the mice with *Stx12* deficiency were lethal, approximately 80% of *Stx12* morpholino-injected zebrafish embryos failed to survive beyond 6 hours post fertilization (hpf), with only about 10% of these embryos could survive for days (Figure 1A), demonstrating that *Stx12* played a vital role in the early development of embryos. Subsequently, the expression pattern of *Stx12* in zebrafish embryogenesis was explored. The representative image of five selected stages (0.2 hpf, 1 hpf, 2 hpf, 3.7 hpf, 6 hpf) were displayed in Figure 1B and quantitative RT-PCR for the embryo stages was performed (Figure 1C), revealing the expression level of *Stx12* peaks within 2-3.7 hours post-fertilization during embryonic development, which led us to choose this particular time window for assessing the impact of *Stx12* knockdown.

**Figure 1.**
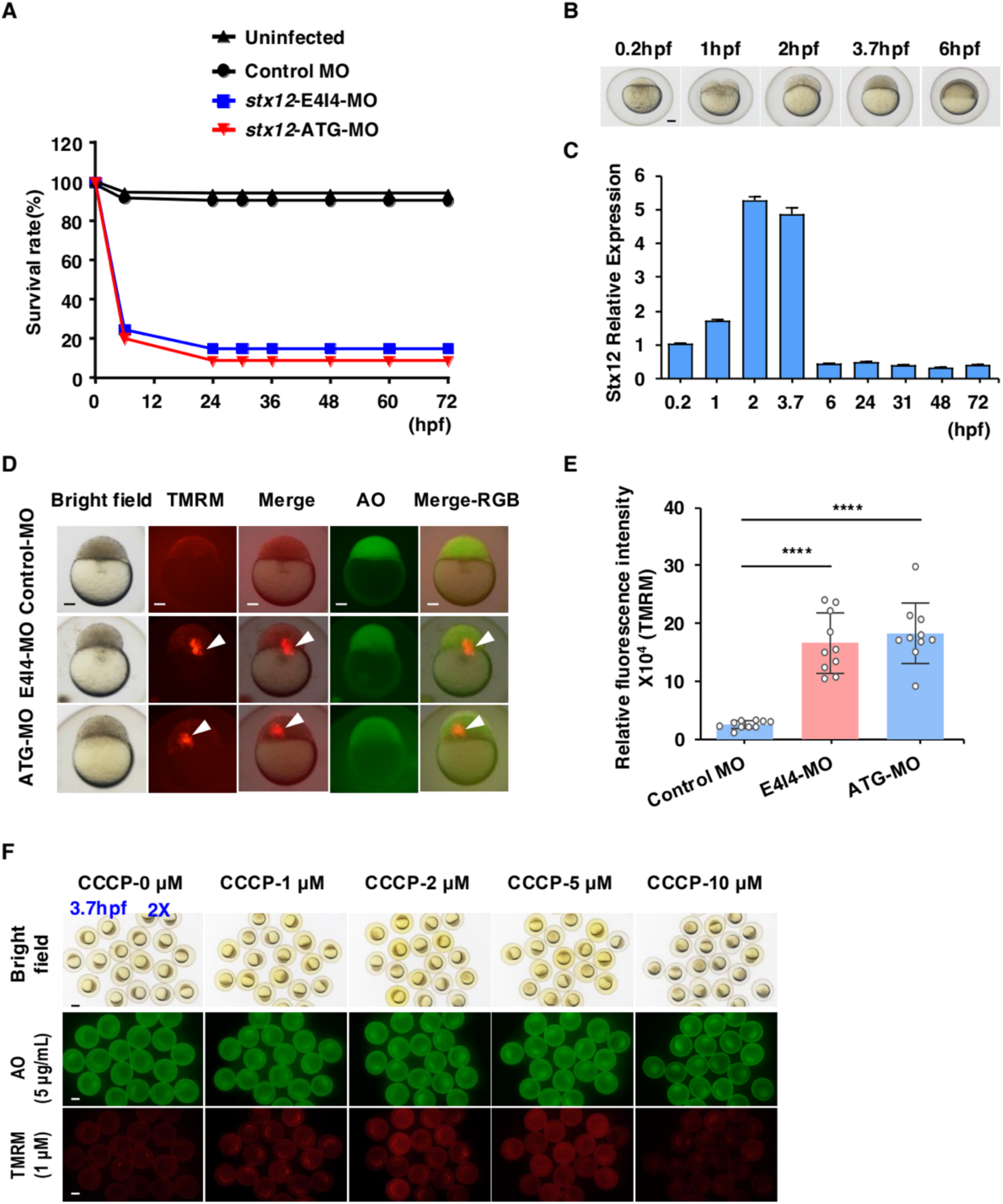
STX12 deficiency induced mitochondrial membrane potential defects in zebrafish. A. A time-course plot of percent survival in control vs. *Stx12* morphants for 3 days. dpf, days post fertilization; hpf, hours post fertilization. B. Representative images of five selected stages of zebrafish embryo. Scale bar: 100 μm. C. qRT-PCR for five embryo development stages (0.2 hpf, 1 hpf, 2 hpf, 3.7 hpf and 6 hpf) demonstrates different expression patterns of *Stx12* during embryonic development. D. Representative images of TMRM staining and AO staining in control, E4I4-MO and ATG-MO zebrafish. Treatment Window: 2 hpf-3.7 hpf; Stage of Image: 3.7 hpf. TMRM Staining: 1 μM; AO Staining: 5 μg/mL. n = 10, Scale bar: 100 μm. E. The quantification of relative TMRM fluorescence intensity. F. Representative images of TMRM staining and AO staining after CCCP treatment. Scale bar: 300 μm

### STX12 deficiency induces mitochondrial defect but not apoptosis in the zebrafish

Apoptosis and mitochondrial dysfunction play an important role in embryogenesis and tissue homeostasis^14,15^. Acridine orange (AO) staining allows us to examine the role of *Stx12* in apoptosis^16,17^ while TMRM (Tetramethylrhodamine methyl ester, perchlorate, biotium) staining enables the investigation of the involvement of STX12 in mitochondrial function^18^. The mitochondrial membrane potential (ΔΨ_m_) was estimated by monitoring fluorescence aggregates of TMRM ^19,20^. To assess the effectiveness of AO staining and TMRM staining in zebrafish embryos, wildtype (WT)-uninjected AB strain zebrafish embryos were exposed to different concentrations of AO (0.25, 0.5, 1.67 and 5 μM) and TMRM (0.05, 0.1, 0.3 and 1 μM) in fish water from 2-hpf to 3.7-hpf at 28.5 ℃ in the dark and the results demonstrated that 5 μg/ml AO and 1 μM TMRM exhibited a good fluorescence staining (Figure S2A). Accordingly, we stained zebrafish embryos with 5 μg/ml AO and 1 μM TMRM in fish water from 2-hpf to 3.7-hpf at 28.5 ℃ in the dark. Here, 1 μM TMRM was used in quenching mode, in which condition the fluorescence intensity was weaker as the probes accumulate in mitochondria at high and quenching concentrations^21^. Intriguingly, there was evidently stronger TMRM fluorescence signal, in the embryos of zebrafish with ATG MO and 4I4 MO (Figure 1D). In contrast, Compared to zebrafish with control-MO, there was no apparent apoptosis in the embryos of zebrafish with ATG MO and 4I4 MO (Figure 1D). The intensity of TMRM fluorescence was quantified in Figure 1E. The increase of TMRM fluorescence intensity was consistent with the effect of 1 μM CCCP (Carbonyl cyanide 3-chlorophenylhydrazone), a proton uncoupler that depolarizes mitochondria, observed in WT-uninjected zebrafish embryos (Figure 1F). This indicated that *Stx12* knockdown in zebrafish prompted mitochondrial depolarization, resulting in a reduction in ΔΨ_m_ with higher TMRM intensity in quenching mode. In addition, toxicity and safety of CCCP in zebrafish model was also explored. The results indicated that 1 μM CCCP did not induce developmental delay or mortality in zebrafish embryos, confirming the safety of the 1 μM CCCP concentration, conversely, 10 μM CCCP led to complete mortality in zebrafish embryos (Figure S2B and S2C).

Collectively, our investigation revealed that the knockdown of *Stx12* results in embryonic lethality and a reduction in ΔΨ_m_ in zebrafish.

### The deficiency of STX12 elicited an inflammation response in mice, with the inflammation being predominantly localized in the lung tissue

Building on the observation of impaired mitochondrial membrane potential in zebrafish embryos, we aimed to explore whether a similar phenotype occurs in a mouse model. To this end, we co-cultured primary mouse embryo fibroblasts from *Stx12*^−/−^ mice and its littermate controls and evaluated mitochondrial membrane potential using 50 nM TMRM staining (unquenching mode) (Figure 2A). The *Stx12^−/−^* MEFs were tagged with an EGFP fluorescent marker, which was incorporated during the gene editing process, serving as an indicator for *Stx12* knockout MEFs. In agreement with the findings in zebrafish, the mitochondrial membrane potential of MEFs from *Stx12^−/−^* mice showed a reduction of mitochondrial membrane potential compared to those from littermate controls, as indicated by decreased TMRM fluorescence intensity (Figure 2B).

**Figure 2.**
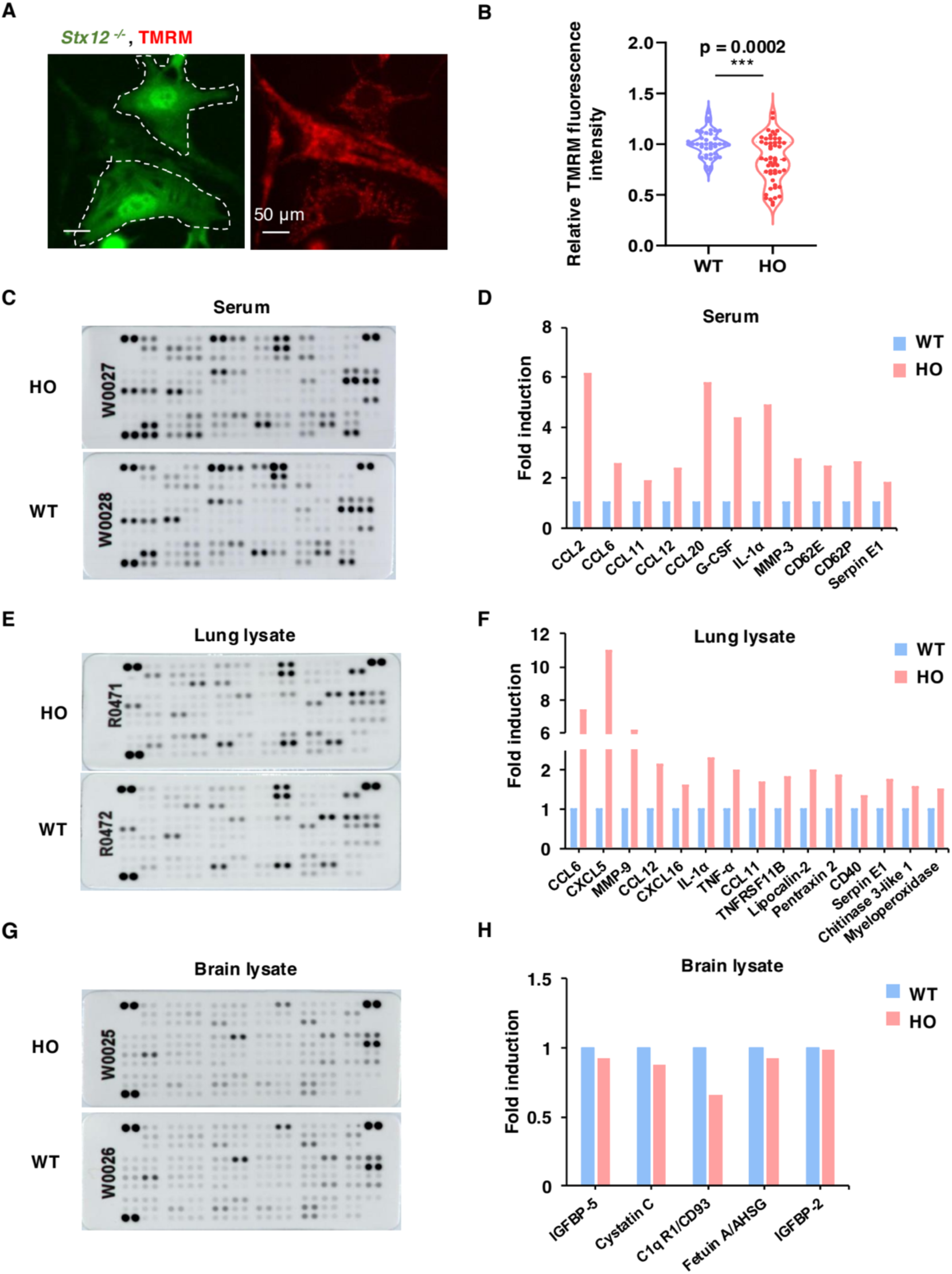
The loss of STX12 elicited an inflammatory response predominantly in the lung tissue. **A.** Representative images of TMRM staining (50 nM) of MEF from *Stx12*^−/−^ mice and littermate controls. **B.** The quantification of relative TMRM fluorescence intensity. Results are presented as mean ± SEM; statistical significance was assessed by Student’s t-test. **C.** GO enrichment bubble plot of differentially upregulated genes in *Stx12*^−/−^ mice lung tissue. The bubble plot illustrates the significantly enriched Gene Ontology (GO) terms for biological processes (BP), molecular function (MF) and cellular component (CC) related to the differentially upregulated genes. The x-axis represents the Gene Ratio while the y-axis lists the corresponding GO terms. The size of each bubble reflects the number of upregulated genes associated with each GO term, and the color of the bubbles denotes the statistical significance (adjusted p-value) of the enrichment. **D, E.** The cytokine kit displays of serum from *Stx12*^−/−^ mice (HO) and the littermate controls (WT, *Stx12*^+/+^) (R&D Systems, Catalog # 410-MT) and quantitative analysis. **F, G.** The cytokine kit displays of lung lysate from *Stx12*^−/−^ mice (HO) and the littermate controls (WT, *Stx12*^+/+^) (R&D Systems, Catalog # 410-MT) and quantitative analysis.

Based on its pale, swollen, and stiff phenotype as demonstrated in our previous study^13^, we speculated that there was inflammation activation in *Stx12*^−/−^ mice. Therefore, we performed a cytokine array (Figure 2C) to evaluate the expression of inflammatory cytokine in the serum of *Stx12*^−/−^ mice (HO) and the littermate controls (WT). The results revealed that *Stx12*^−/−^ mice exhibit significant elevated levels of several inflammatory cytokines, including chemokines (CCL2, CCL6, CCL11, CCL12, CCL20), cytokine (G-CSF, IL-1α), adhesion molecules (CD62E, CD62P), Serpin E1, and MMP-3, thus confirming our initial hypothesis (Figure 2D). To further investigate the tissue-specific origins of inflammation, we assessed cytokine expression levels in homogenates from multiple tissues including lung (Figure 2E), brain (Figure 2G), spleen (Figure S3A), liver (Figure S3C) and kidney (Figure S3E). The statistical results unveiled a significant activation of inflammation in lung tissue (Figure 2F), no cytokine elevation in the brain lysate (Figure 2H) and minor increases in the lysates of spleen (Figure S3B), liver (Figure S3D), and kidney (Figure S3F). The results highlighted that the lung was the primary site of inflammation, guiding our research focus accordingly.

Next, we performed bulk RNA sequencing (RNA-seq) of lung tissues from *Stx12*^−/−^ mice and their littermate controls in E19.5. The gene expression profiles between the HO (*Stx12*^−/−^) and WT groups identified 13,340 co-expressed genes, 597 genes specifically expressed in the HO group, and 273 genes specifically expressed in the WT group (Figure S4A). The differentially expressed genes between the two groups were also analysed. We defined differentially expressed genes with an absolute log-fold change above 1 and an FDR-adjusted p-value of less than 0.05, yielding differences in 901genes between HO and WT mice with 806 genes upregulated and 95 genes downregulated and the volcano plot was shown in Figure S4B. GO enrichment analysis of the differentially upregulated genes revealed several significantly enriched terms within the Biological Process (BP), Molecular Function (MF), and Cellular Component (CC) categories, as illustrated in the bubble plot (Figure S4C). The prominent categories were primarily associated with the regulation of inflammatory response, leukocyte migration and cytokine activity, consistent with the inflammation activation observed in the cytokine array of lung lysate from *Stx12*^−/−^ mice.

### The deficiency of STX12 triggered mitochondrial damage and mitochondrial DNA release into cytosol in alveolar type II epithelial cells

Building on our findings of mitochondrial dysfunction in STX12-knockdown zebrafish embryos and STX12-knockout mouse MEFs, we further explored the effects of STX12 deficiency on mitochondrial function of lung tissue. Considering that mitochondria generate most of the cell’s energy through oxidative phosphorylation (OXPHOS), we examined the expression levels of OXPHOS subunits in the lung tissue of *Stx12^−/−^* mice and their littermate controls to evaluate mitochondrial function using western blotting assay, with quantification conducted via Image J (Figure 3A, B). Our analysis revealed a significant reduction in the expression levels of mitochondrial complex subunits I, II, and V in *Stx12^−/−^* mice compared to the ones of their littermate controls, while the levels of subunits III and IV showed no significant change. Meanwhile, Gene Set Enrichment Analysis (GSEA) on RNA-seq data from *Stx12^−/−^* lung mice revealed that the most prevalent altered biological processes were associated with pathways linked to inflammation and the innate immune response. Additionally, there was a notable enrichment of the cytosolic DNA-sensing pathway (Figure 3C). Next, we cultured primary alveolar type II epithelial cells (AEC II) and performed immunofluorescence staining with antibody specific to DNA and TOM20 to detect cytosolic DNA. Statistical analyses demonstrated a marked increase in the number of cytosolic DNA foci per cell, as well as a higher proportion of cells containing cytosolic DNA foci in AEC II isolated from *Stx12^−/−^* mice, compared to those from littermate controls (Figure 3D, E, F). These findings suggest a significant accumulation of cytosolic DNA in the alveolar type II epithelial cells of *Stx12^−/−^* mice. Previous studies have demonstrated the involvement of mitochondrial DNA (mtDNA) and mitochondrial RNA (mtRNA) as potent triggers of nucleic acid sensing pathways, resulting in the activation of different inflammatory pathways^22^. In light of the mitochondrial dysfunction observed, we hypothesized that the cytosolic DNA foci might originate from the release of mitochondrial DNA (mtDNA). To validate this, we quantified the copy number of mitochondrial DNA (mtDNA) in isolated cytosolic fractions of alveolar type II epithelial cells (AEC II) using quantitative polymerase chain reaction (qPCR) (Figure 3G). In line with our hypothesis, transcript levels of mitochondrial genes mt-ND2, mt-CyTb and mt-Dloop1 were significantly elevated in the cytosolic compartments of *Stx12^−/−^* alveolar type II epithelial cells, in comparison to the corresponding littermate controls.

**Figure 3.**
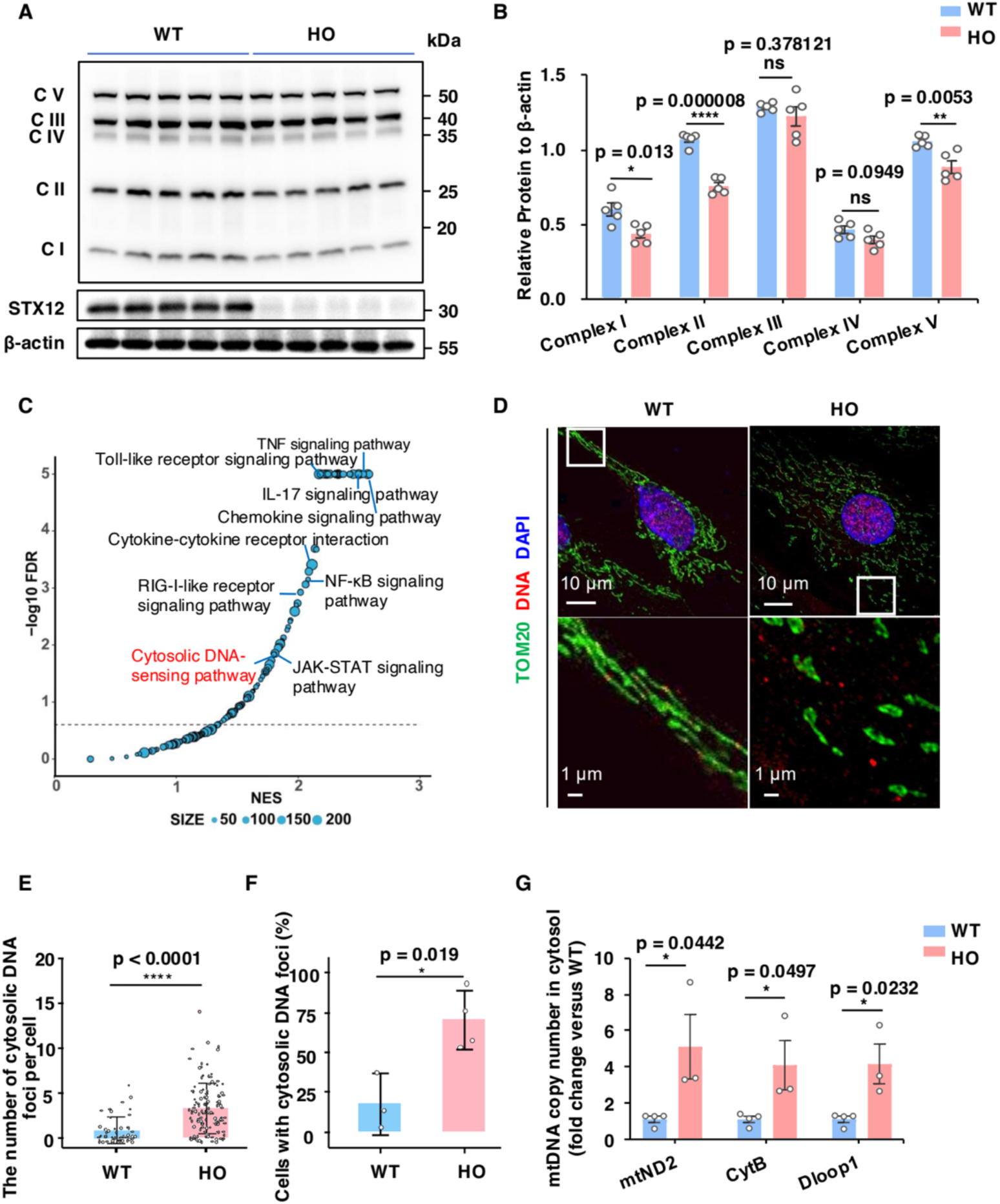
The ablation of STX12 triggered mitochondrial damage and cytosolic release of mitochondrial DNA in alveolar type II epithelial cells. **A.** SDS-PAGE of mitochondrial OXPHOS subunits in lung lysates using OXPHOS cocktail antibodies. **B.** Quantification of the levels of C I (complex I), C II (complex II), C III (complex III), C IV (complex IV) and C V (complex V) in Figure 3A. Results are presented as mean ± SEM; statistical significance was assessed by Student’s t-test. **C.** Volcano plots of the GSEA, highlighting the differentially regulated pathways in *Stx12*^−/−^ mice lung tissue. FDR, false discovery rate; NES, normalized enrichment score. **D.** Representative confocal images of immunofluorescence of primary alveolar type II epithelial cells (AEC II cells) using antibodies against TOMM20 (mitochondria) and DNA. (WT n = 4; KO n = 3 independent cultures) Boxes mark the enlarged images shown below. **E, F.** Number of cytosolic DNA foci per cell (E) and percentage of AEC II cells showing cytosolic DNA foci (F) from D. WT n = 3 HO n = 4 independent experiments. Results are presented as mean ± SD; statistical significance was assessed by Student’s t-test. G. Quantification of mtDNA copy number by qPCR using mtND2, CytB and Dloop1 primers, from isolated cytosolic fractions of AEC II cells of WT and *Stx12*^−/−^ mice. Results are presented as mean ± SEM; statistical significance was assessed by Student’s t-test.

These findings demonstrated that the knockout of *Stx12* resulted in impaired mitochondrial oxidative phosphorylation and the release of mitochondrial DNA into the cytoplasm, which might account for the activation of lung inflammation observed in the *Stx12^−/−^* mice.

### Loss of *Stx12* elicits the activation of cGAS-STING-TBK1 pathway and Type I interferon pathway

In addition to detecting cytoplasmic DNA foci at the cellular level, these foci were also observed at the tissue level within the lung. 30-micron frozen lung tissue sections obtained from *Stx12^−/−^*mice and littermate controls were stained with antibodies specific to DNA and TOM20, followed by imaging under fluorescence microscopy (Figure 4A). The DNA foci that did not colocalize with TOM20 and DAPI were identified as cytoplasmic DNA. These foci were quantified and subjected to statistical analyses. Comparative results revealed that *Stx12^−/−^* mice exhibited a significant increase in the number of cytoplasmic DNA foci per cell and per field compared to the corresponding littermate controls (Figure 4B, C).

**Figure 4.**
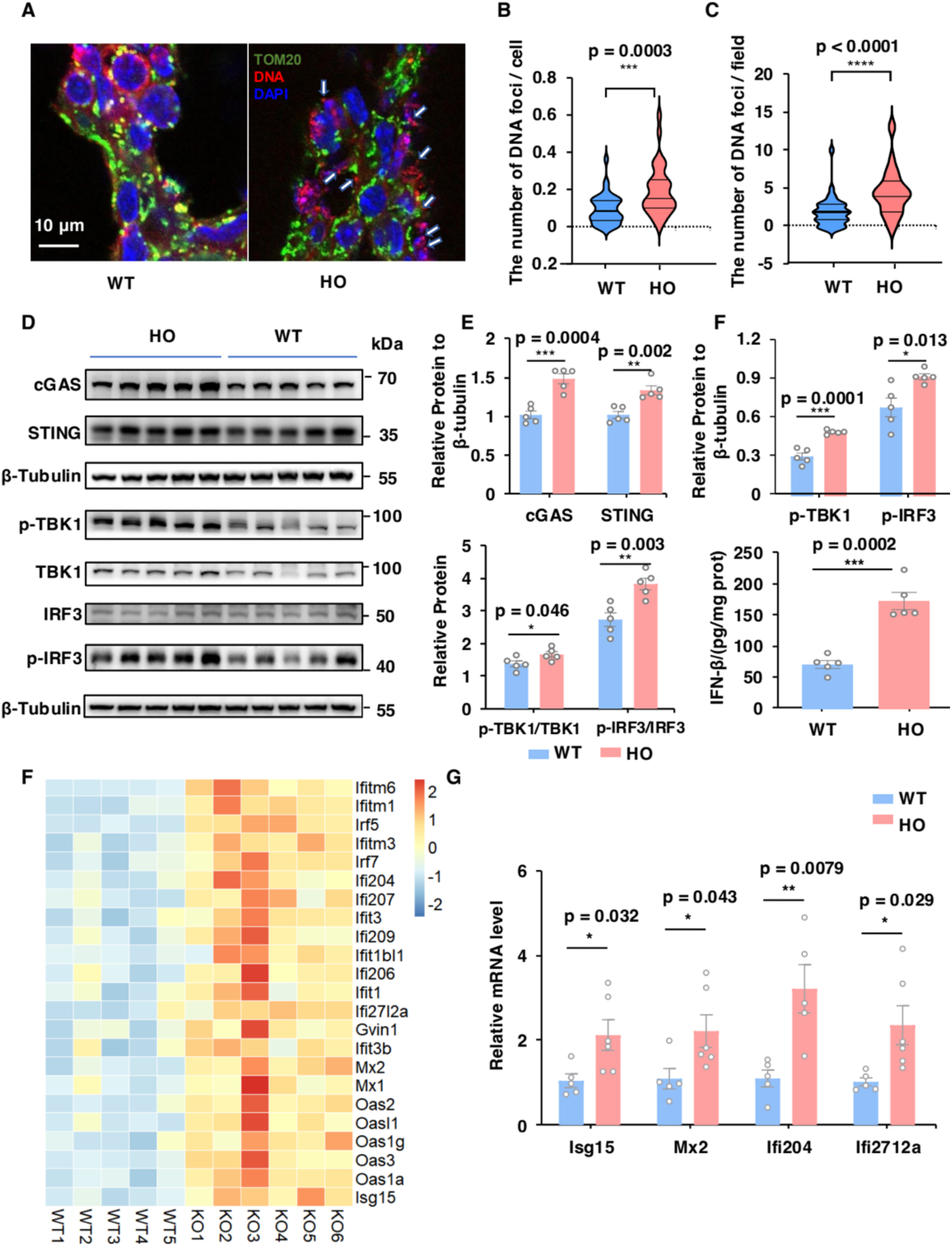
The cGAS–STING pathway and Type I interferon pathway were activated of in lung tissue of *Stx12*^−/−^ mice. **A.** Representative confocal images of immunofluorescence of lung frozen slice (30 μm) using antibodies against TOMM20 (mitochondria) and DNA. White arrowheads indicate cytosolic DNA foci. Scale bar, 10 μm. **B.** Quantification of the average cytosolic DNA puncta per cell in the corresponding field. WT n = 3, KO n = 4 independent experiments. **C.** Quantification of the average cytosolic DNA puncta per field. WT n = 3, HO n = 4 independent experiments. **D.** The expression of cGAS, STING, the phosphorylation of TBK1 (p-TBK1), the phosphorylation of IRF3 (p-IRF3) and GAPDH by western blot in lung lysate from *Stx12*^−/−^ and the littermate control (WT, *Stx12*^+/+^) mice. **E.** Serum IFN-β levels from *Stx12*^−/−^ and the littermate control (WT, *Stx12*^+/+^) mice (n = 5 per group) by enzyme-linked immunosorbent assay. Results are presented as mean ± SEM; statistical significance was assessed by Student’s t-test. **F.** Heat map showing upregulated interferon stimulates genes (ISG) based on RNA-seq data of *Stx12*^−/−^ lung tissue. **G.** Expression levels of ISGs by qRT-PCR from the lung tissue of *Stx12*^−/−^ (n = 6) and the littermate control (WT, *Stx12*^+/+^) (n = 5) mice. The mRNA expressions of target genes were normalized to β-actin. Results are presented as mean ± SEM; statistical significance was assessed by Student’s t-test.

Previous extensive research has demonstrated cGAS-STING pathway (cyclic GMP-AMP synthase-stimulator of interferon genes) is a crucial component of the innate immune system, responsible for detecting cytosolic DNA from various sources, including microbial pathogens, cancer cells, or DNA damage, and initiating an immune response^23,24,25^. Therefore, we hypothesized that the activation of the cGAS-STING pathway might play a role in the observed phenotypes of *Stx12* knockout mice. The results of western blotting revealed that *Stx12* knockout markedly increased the level of cGAS and STING, and dramatically elevated the phosphorylation of TBK1 and IRF3 in mouse lung lysate (Figure 4D-G), suggesting the activation of cGAS-STING signaling pathway in the lung of *Stx12^−/−^*mice. Furthermore, enzyme-linked immunosorbent assay (ELISA) analysis revealed a statistically significant elevation in the level of IFN-β within the lung tissue of *Stx12^−/−^* mice compared to that of control subjects (Figure 4H). Moreover, transcriptomic analysis of lung tissue demonstrated that a substantial proportion of upregulated transcripts in *Stx12^−/−^* mice were encoded by interferon-stimulated genes (ISGs), as depicted in Figure 4I. To confirm the upregulation of these ISGs expression including genes such as Isg15, Mx2, Ifi204 and Ifi2712a in *Stx12^−/−^*mice, reverse transcription quantitative polymerase chain reaction (RT–qPCR) was further used to validate (Figure 4J). Our findings illustrated the sequential activation of interferon signaling pathway following the engagement of cGAS-STING-pTBK1-pIRF3 cascade, providing a comprehensive understanding of the inflammatory response in *Stx12^−/−^* mice.

In summary, cGAS-STING pathway and the Type I interferon signaling pathway were markedly activated by the presence of cytoplasmic mtDNA in the lung tissue of *Stx12^−/−^*mice.

### Neutrophils-related severe immune response in *Stx12* knockout mice

RNAseq analysis was further performed to comprehensively elucidate cellular pathways affected by STX12 deficiency. KEGG pathway enrichment analysis on the differentially upregulated genes revealed significant enrichment in inflammation-related pathways and the pathways with −log10 (padj) > 5.5 were displayed in Figure 5A. Notably, the IL-17 signaling pathway and neutrophil extracellular trap (NET) formation pathway were prominently enriched, along with GO enrichment of neutrophil migration and neutrophil chemotaxis (Figure S4C), suggesting a crucial role of neutrophils in the pulmonary inflammatory response of *Stx12^−/−^*mice. Consistent with these findings, the blood smear results revealed a marked increase in the number of neutrophils (Figure 5B) in *Stx12^−/−^* mice compared to *Stx12^+/+^* littermate controls, with no significant changes observed in lymphocyte and monocyte (Figure S5A and S5B). Meanwhile, qPCR analyses revealed upregulated expression of neutrophil markers, including Ly6g, Myeloperoxidase (MPO), ELANE and CSF3R, as well as elevated levels of CSF3, C3, CXCL1, CXCL2, CXCL5 and TNF, which served as key chemokines for neutrophil recruitment and activation in the lung tissue of *Stx12^−/−^* mice compared to littermate controls (Figure 5C). S100A9, primarily secreted by neutrophils and can promote the migration and activation of neutrophils^26^, was also found to be significantly upregulated and confirmed by western blot analysis (Figure 5F and 5G). These observations indicated a pronounced neutrophil infiltration in the lung tissue of *Stx12^−/−^* mice. In addition, *Stx12^−/−^* lung homogenate showed increased IL-1*β* (Figure 5D), which can activate neutrophils^27^ and promote the formation of NETs^28^. The heatmap indicated an accumulation of various cytokines in the lung tissue of *Stx12^−/−^* mice, including neutrophil-related elements, interleukins, TNFs-related, complement components-related, colony-stimulating factors, and chemokines (Figure 5H), with some further validated by qPCR (Figure S5C). Additionally, the upregulation of IL-6 was confirmed through qPCR (Figure 5E) and western blot analysis (Figures 5E-G). These cytokines and granule enzymes (MPO, Elane, MMP) further attracted and activated neutrophils, creating a robust and cascading inflammatory response within the lung tissue of *Stx12^−/−^* mice. We hypothesized that this might be the cause of mortality in *Stx12* knockout mice. To investigate this, we also conducted rescue experiments using various interventions on pregnant mice, including the immunosuppressant rapamycin (intravenous injection; once a day; 1.6 mg/kg), the neutrophil elastase inhibitor Sivelestat (intravenous injection; once a day; 50 mg/kg, DMSO 5%), immunomodulators Vitamin C and Vitamin D^29,30^ (via gavage; Vc: 205 mg/kg in double distilled water; Vd: 150 µg/kg in double distilled water, final ethanol < 1/10000), VDAC inhibitor VBIT-4 ^31^ (provided in the drinking water; 20 mg/kg, final pH 5.0, DMSO 0.05%) and mitochondrial stabilizer Taurine (provided in the drinking water; 1000 mg/kg/day in double distilled water). Given the lethality of STX12 knock out mice at birth, it is not feasible to administer these interventions directly to the knockout mice. However, none of these interventions succeeded in rescuing the phenotype.

**Fig 5.**
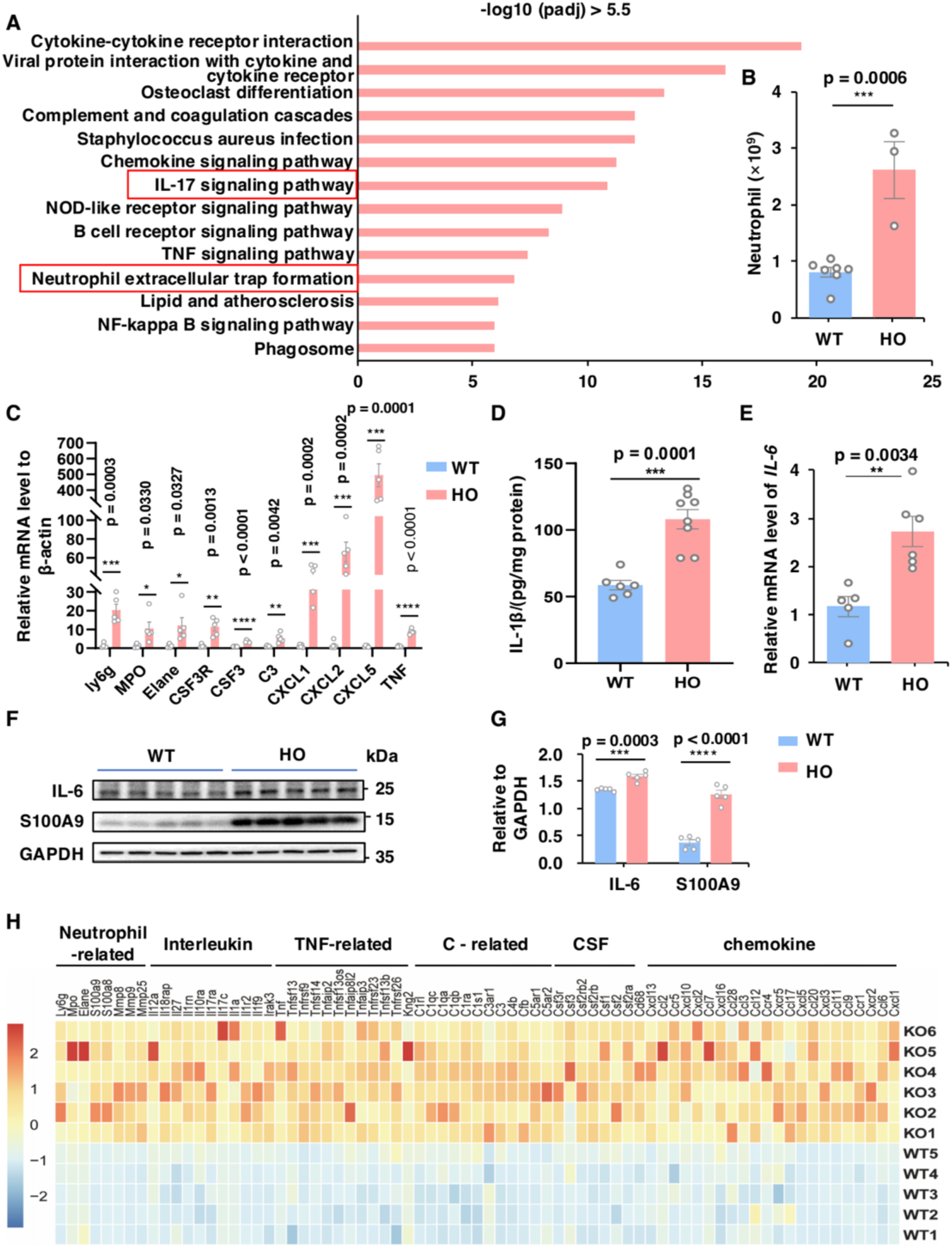
Neutrophils are involved in the severe inflammation in the lung tissue of *Stx12*^−/−^ mice. **A.** KEGG pathway analysis based on differentially upregulated expressed genes in *Stx12*^−/−^ mice (-log_10_(padj) > 5.5) from RNA-seq data. **B.** The number of neutrophil in the blood from *Stx12*^−/−^ and the littermate control (WT, *Stx12*^+/+^ mice. **C.** The relative mRNA level of neutrophil markers (Ly6g, MPO, ELANE, and CSF3R) and key chemokines for neutrophil recruitment and activation (CSF3, C3, CXCL1, CXCL2, CXCL5, and TNF) normalized to β-actin. WT n = 5, HO n = 5 independent samples. **D.** IL-1β levels in the lung lysate from *Stx12*^−/−^ (n = 8) and the littermate control (WT) mice (n = 6) by enzyme-linked immunosorbent assay. **E.** Relative mRNA levels of IL-6 from lung of *Stx12*^−/−^ (HO) and the littermate control (WT, *Stx12*^+/+^) mice. **F.** The expression of IL-6, S100A9 and GAPDH by western blot in lung lysate from *Stx12*^−/−^ (HO) and the littermate control (WT, *Stx12*^+/+^) mice. **G.** Quantification of the levels of IL-6 and S100A9 normalized to GAPDH. **H.** Heatmap showing upregulated inflammation-related gene including neutrophil-related gene, interleukin, tumor necrosis factor (TNF)-related gene, complement(C) - related gene, colony stimulating factor (CSF)-related gene and chemokines in WT and *Stx12*^−/−^ lung tissue. **I.** The picture of newborn mice after drug intervention. Results are presented as mean ± SEM; statistical significance was assessed by Student’s t-test.

**Figure 6.**
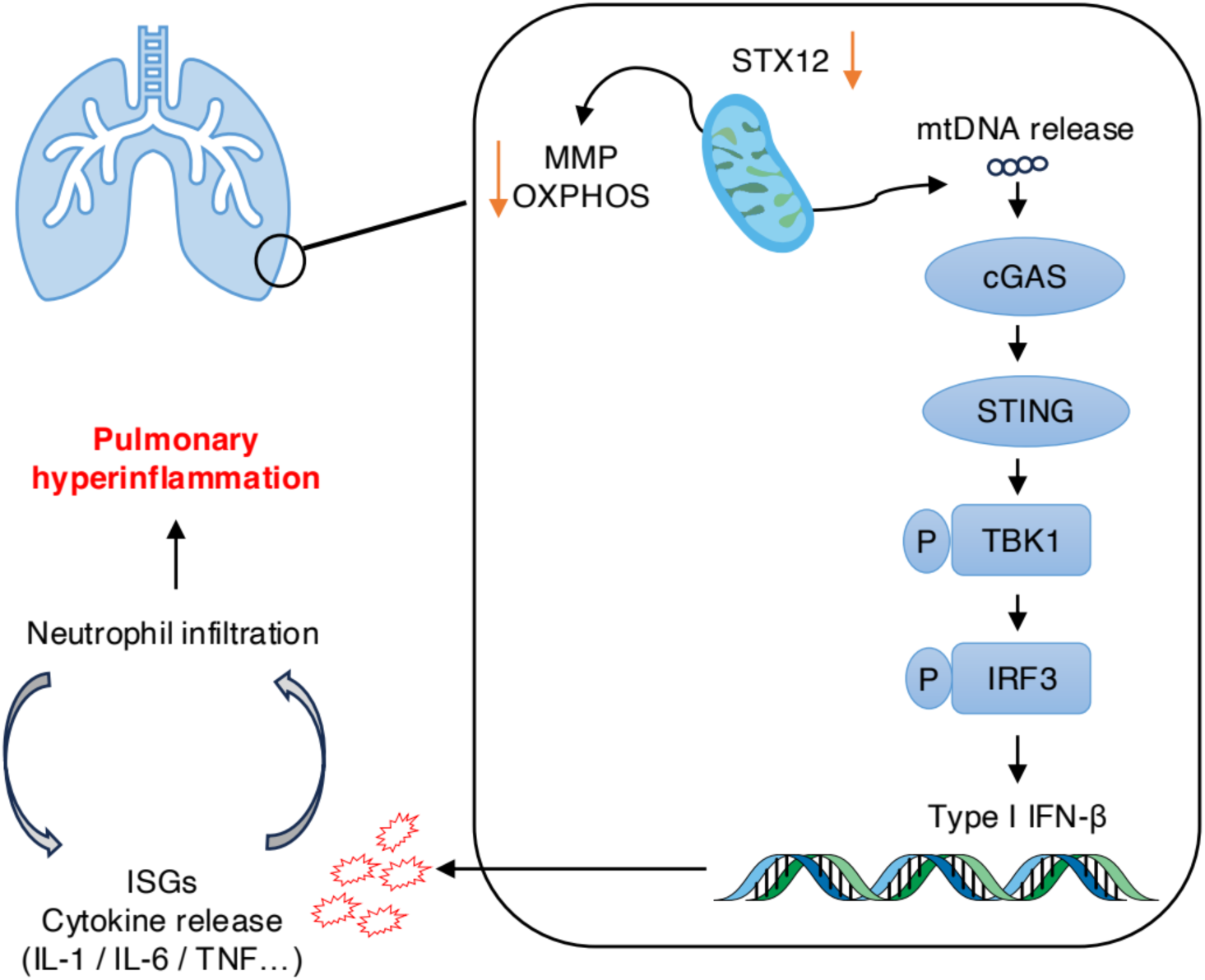
Schematic of systemic immune-inflammation in STX12-KO mice. The ablation of STX12 leads to decreased mitochondrial membrane potential (MMP), reduced expression levels of mitochondrial complex subunits, and the release of mitochondrial DNA (mtDNA). Then, mtDNA release activates the cGAS-STING-pTBK1-pIRF3 pathway, subsequently triggering Type I interferon response and downstream interferon-stimulated genes (ISGs) and cytokines in lung tissue. Additionally, cytokines release and neutrophil infiltration mutually enhance each other, forming an amplified cascade of hyperinflammation, referred to as “cytokine storm”, which potentially contribute to the mortality observed in *Stx12* knockout mice.

## Discussion

SNAREs constitute a critical family of proteins that facilitate the intricate process of membrane fusion within eukaryotic cells^32^. SNARE proteins are characterized by their ability to form a complex that brings the membranes of vesicles (v-SNARE) and target compartments (t-SNARE) into close proximity, facilitating the fusion process^33,34,35,36^. SNARE proteins are integral to numerous fundamental processes, including the initiation and extension of neurites, the specification and elongation of axons, as well as synaptogenesis and synaptic transmission^37,38^ and are critically implicated in a range of diseases such as neurodegenerative disorders including Alzheimer’s disease (AD)^39,40,41,42,43^, Parkinson’s disease (PD), Amyotrophic Lateral Sclerosis (ALS), Huntington’s disease (HD) and Spinal Muscular Atrophy (SMA)^38^, metabolic conditions^44^ and cancer^45,46^.

As for the roles of SNARE proteins in mitochondrial dynamics, some evidence suggests that STX17 promotes autophagy formation by localizing at the contact sites between the endoplasmic reticulum and mitochondria^47^ and is integral to the process of autophagosome maturation by mediating autophagosome-lysosome fusion and thus facilitating cellular homeostasis^48^. Syntaxin 4 (STX4), when enriched in skeletal muscle, reverses peripheral insulin resistance and ameliorates mitochondrial dynamics via the regulation of the dynamin-related protein 1 (Drp1)^49^ and plays a pivotal role in the lysosome-associated exocytosis of mitochondria, which is independent of mitophagy and contributes to the parkinsonism-like symptoms induced by flunarizine (FNZ) ^50^. STX5 facilitates cholesterol trafficking from plasma membranes to mitochondria for adrenal steroid synthesis^51^. Furthermore, existing research has demonstrated that STX17 mitigates heart failure by promoting Drp1-dependent mitophagy via recruitment of CDK1^52^ and plays a role in obesity cardiomyopathy through facilitating obesity-induced mitochondria-associated endoplasmic reticulum membranes (MAMs) formation and mitochondrial Ca^2+^ overload^53^. Here, we showed that STX12 depletion induced decreased mitochondrial membrane potential in zebrafish (Figure 1D) and mouse embryo fibroblasts (Figure 2A and 2B). Moreover, *Stx12* knockdown leads to reduced expression levels of mitochondrial complex subunits and mtDNA release in mouse lung tissue, activating the cGAS-STING signaling pathway, ultimately leading to pulmonary inflammation in mice.

Transcriptome and inflammation analyses reveal the enrichment of cytokines and chemokines in *Stx12^−/−^* lung tissue. Gene enrichment analysis implies a critical role of IL-17-neutrophil axis in the inflammatory process. The enriched IL-17 signaling pathway has been implicated to intimately involve in the regulation of neutrophil function encompassing migration, activation, and degranulation of these cells^54^. NETs are complex structures composed of DNA, histones, and antimicrobial proteins that are expelled by neutrophils as a critical component of the immune response to ensnare and neutralize a broad spectrum of pathogens^55^. The formation of NETs is a highly regulated process that can be triggered by various stimuli, including IL-17. Once activated, neutrophils undergo a unique form of cell death that leads to the release of NETs, which can effectively immobilize and kill a wide range of microorganisms^56^. However, excessive or uncontrolled NET formation can also contribute to tissue damage and has been implicated in the pathogenesis of several inflammatory diseases, including acute lung injury and acute respiratory distress syndrome ^57^. The substantial enrichment of the IL-17 signaling pathway and NETs formation pathway in the KEGG analysis underscores the pivotal role of neutrophils in the pulmonary inflammation observed in *Stx12* knockout mice (Figure 5A). Furthermore, the GO analysis, which emphasizes the chemotaxis and migration of leukocytes—especially neutrophils—provides additional support for this conclusion (Figure S4C). For instance, ly6G is a glycoprotein that is predominantly expressed on the surface of mouse neutrophils and is commonly used as a marker of neutrophil ^58,59^. MPO is a key enzyme found in the azurophilic granules of neutrophils and plays a crucial role in the antimicrobial defense of neutrophils by producing hypochlorous acid^60,61^. Particularly, MPO has been implicated in the regulation of various neutrophil functions, including neutrophil activation^62,63^, trafficking^63,64^, phagocytosis^63^, lifespan^65^ and formation of extracellular traps^61^ and MPO-triggered autoimmunity^63,66^. MPO functions as a survival signal for neutrophils and thereby contribute to prolongation of inflammation^65^. Elane, also known as Neutrophil Elastase, is a serine protease enzyme found in the azurophilic granules of neutrophils. Elane has been shown to degrade almost all extracellular matrix proteins^67^, disrupt the structural integrity of the tissue microenvironment^68^, and induce the release of pro-inflammatory cytokines such as interleukin-6, interleukin-8 and so on^69,70^. These significant elevations of Ly6G, MPO and Elane in the *Stx12^−/−^* lung tissue indicate substantial neutrophil infiltration and potential damage of lung tissue. These injuries further exacerbate the release of cytokines and chemokines. S100A9, highly present in the cytoplasmic fraction of neutrophils, is associated with increased production of inflammatory cytokines^71^ and can promote the migration and activation of neutrophils^26^. Besides, MMPs released by activated neutrophils have been demonstrated to modulate inflammatory responses by influencing the levels and activities of cytokines and chemokines, including the cleavage of several CXC chemokines in addition to degrading the extracellular matrix and facilitating neutrophil migration^72^. Furthermore, the increased cytokines such as IL-1, IL-6, TNF, CSF, C3 and neutrophil-specific chemokines, particularly CXCL1, CXCL2, and CXCL5 could also promote the chemotaxis and activation of neutrophils^73,74,75,76,77^. Therefore, we proposed that the infiltration of neutrophils and the release of cytokines in the lung tissue mutually enhance each other, creating a cascade amplification of inflammation, commonly referred to as “cytokine storm.”^78,79^ Consequently, we employed various interventions, but none effectively mitigated the lethal phenotype. On the one hand, it is conceivable that the placental barrier significantly diminishes the fetuses’ absorption of these drugs, which may account for the lack of efficacy observed. The permeability of VBIT-4 through the placental barrier, in particular, remains uncertain. On the other hand, the observed lethality in the mice might result from a combination of systemic effects including multi-system failure, anemia, inflammation, and other potential complications. These findings suggest that the underlying mechanisms are complex and multifactorial, necessitating further detailed investigation to elucidate the precise causes of mortality in *Stx12^−/−^*mice.

Overall, the cGAS-STING activation trigged by pulmonary mtDNA release in *Stx12^−/−^*mice is a new mechanism of pulmonary inflammation, expanding our understanding of how mitochondria could affect innate immune responses and providing a basis for further investigations of mitochondria injury-driven immunopathology.

## Material and methods

### Zebrafish Care and Maintenance

Adult wild-type AB strain zebrafish were maintained at 28.5 ℃ on a 14 h light/10 h dark cycle. Five to six pairs of zebrafish were set up for nature mating every time. Embryos were maintained at 28.5 ℃ in fish water (0.2% Instant Ocean Salt in deionized water).

### Zebrafish microinjections

The morpholino (MO) were designed and synthesized by Gene Tools, LLC (http://www.gene-tools.com/). Antisense MOs (GeneTools) were microinjected into fertilized one-cell stage embryos according to standard protocols.

*Stx12* ATG-MO 5′- TGGAGCAAACTACAGCAGGAAGCCA -3′

*Stx12* E4I4-MO 5′- ACTGGCAACTACAAAAGTACCTGTT -3′

Control-MO 5’- CCTCTTACCTCAGTTACAATTTATA -3’

The dose of the MOs used for injection was as follows: Control-MO and E4I4-MO, 4 ng per embryo; ATG-MO, 4 ng per embryo. Primers spanning *Stx12* exon 2 and exon3 (forward primer: 5‘- CACACTGAATACCGCTCAAATC -3’) and exon 5 (reverse primer: 5‘- CCACTGACTCCTTCTCTTTCTC -3’) were used for RT-PCR analysis for confirmation of the efficacy of the E4I4-MO.

### Acridine orange staining for apoptosis

Control-MO injected embryos and embryos injected with *Stx12*-MO were immersed in 5 μg/ml AO (acridinium chloride hemi-[zinc chloride], Sigma-Aldrich) in fish water from 2-hpf to 3.7-hpf at 28.5 ℃. Next, zebrafish embryos were rinsed thoroughly in fish water three times (5 min/wash) and then oriented on their lateral side and mounted with methylcellulose in a depression slide for observation by fluorescence microscopy.

### TMRM staining in zebrafish

The mitochondrial membrane potential (ΔΨ_m_) was estimated by monitoring the fluorescent intensity of TMRM (Tetramethylrhodamine methyl ester, perchlorate, biotium)^19,20^. To determine the efficacy of TMRM staining in zebrafish embryos, uninjected WT AB zebrafish embryos were immersed in four different concentrations of TMRM (0.05, 0.1, 0.3, and 1μM) in fish water from 2-hpf to 3.7-hpf at 28.5 ℃ in the dark. After treatment, check the embryos using Nikon SMZ18 fluorescence microscope. The results showed that 1μM TMRM displayed a good fluorescent signal. Control-MO injected embryos and embryos injected with *Stx12*-MO were immersed in 1 μM TMRM in fish water from 2-hpf to 3.7-hpf at 28.5 ℃ in the dark. Next, zebrafish embryos were rinsed thoroughly in fish water three times (5 min/wash) at 3.7 hpf. Zebrafish embryos were then oriented on their lateral side and mounted with methylcellulose in a depression slide for observation by fluorescence microscopy.

### Primary culture of mouse embryo fibroblasts (MEF)

Sacrifice the anesthetized pregnant mouse at embryonic 17.5, then carefully dissect out the uterus. Immerse the uterus into 75% (v/v) ethanol followed by rinsing with PBS without Ca^2+^ and Mg^2+^ (Gibco, Invitrogen). Separate each embryo from its placenta, remove head, limbs and viscera and mince the remaining tissue into pieces as small as possible. Use part of tail for genotype identification. Digest the tissues in fresh 15 ml 0.05% trypsin at 37 °C for 30 min, and shake the 50 ml tube every 10 min. Terminate the trypsin activity by adding an equal volume of MEF medium (10 ml FBS in 90 ml DMEM). Centrifuge the cells at 1000 g for 5 min, discard the supernatant and repeat this step once. Finally, resuspend the cells in 10 ml of MEF medium and culture them in a 55 cm² Petri dish at 37°C with 5% CO2.

### Cytokine array

Collect blood samples from *Stx12* knockout (KO) and wild-type (WT) littermate control mice via decapitation using capillary tubes and allow blood samples to clot at room temperature for 2 hours. Then, centrifuged the samples at 2,000 x g for 15 minutes at 4 °C to obtain serum. Collect tissue samples from *Stx12* knockout (KO) mice and littermate wild-type (WT) controls, and then homogenize tissues in lysis buffer (1% Triton X-100, 20 mM Tris-HCl pH 7.5, 150 mM NaCl, 1% deoxycholate, protease inhibitor cocktail) and centrifuge at 12,000 g for 15 minutes at 4°C. Supernatants were collected, and protein concentrations were determined using the BCA Protein Assay Kit (Epizyme Biomedical). The cytokine profile of the tissue homogenates was assessed using the Mouse Cytokine Array Panel A (R&D Systems) according to the manufacturer’s instructions. Briefly, array membranes were blocked with the blocking buffer provided in the kit for 1 hour at room temperature. Subsequently, 150 µg of tissue protein from each sample or 100 µl serum was incubated with the array membranes overnight at 4°C on a rocking platform. After washing, the membranes were incubated with a cocktail of biotinylated detection antibodies for 1 hour at room temperature. Membranes were then washed again and incubated with streptavidin-HRP for 30 minutes. Chemiluminescent detection was performed using the prepared Chemi Reagent Mix in the kit and the signals were captured using a Chemiluminescence imager (MiNiChemi). Densitometry analysis was conducted using ImageJ software (NIH). The intensity of each cytokine spot was normalized to the positive controls on the membrane, and the relative cytokine levels were quantified to WT.

### Western blotting

Protein extraction was performed using RIPA buffer (Beyotime) supplemented with protease and phosphatase inhibitors (Roche). The protein concentration was determined using the BCA Protein Assay Kit (Epizyme Biomedical). Equal amounts of protein (20 μg) were separated by SDS-PAGE on polyacrylamide gel and transferred to a PVDF membrane (Millipore). The membranes were blocked with 5% BSA in TBS-T (20 mM Tris-HCl, 150 mM NaCl, 0.1% Tween-20, pH 7.6) for 2 hrs at room temperature and incubated overnight at 4°C with primary antibodies diluted in antibody dilution buffer (Beyotime). After washing three times with TBS-T, the membranes were incubated with HRP-conjugated secondary antibodies (1:5000, Proteintech) for 2 hrs at room temperature and detected with chemiluminescence detection system (Millipore, Billlerica, MA). The primary antibodies used included the following: rabbit anti-STX12 monoclonal antibody (1:1000, Custom-made), mouse anti-β-actin (1:5000; Proteintech, #60008-1-Ig), mouse anti-GAPDH (1:20000, Proteintech, #60004-1-Ig), anti-OXPHOS antibody (1:1000; Abcam, #ab110413), anti-cGAS (1:1000; Cell Signaling Technology, #31659), anti-TBK1(1:1000, Cell Signaling Technology, #3504S), anti-pTBK1(1:1000; Cell Signaling Technology, # 5483S), anti-IRF3(1:1000; Proteintech, #11312-1-AP), anti-pIRF3(1:1000; Cell Signaling Technology, #29047), anti-IL-6 (1:1000; Abcam, #ab229381), anti-S100A9 (1:1000; Boster, #PB0718).

### RNA Extraction and RT-qPCR

RNA samples were extracted using Trizol reagent (Invitrogen) and RNA was converted to complementary deoxyribonucleic acid (cDNA) using RevertAid First Strand cDNA Synthesis Kit following the manufacturer’s instructions (Invitrogen). Quantitative reverse transcription-PCR (qRT-PCR) was performed using SYBR Green Master Mix (Takara, Otsu, Shiga, Japan) on the qTOWER³ Series (analytikjena, Germany). Gene expression was calculated using the CT method relative to the housekeeping gene glyceraldehyde-3-phosphate dehydrogenase (GAPDH). The primers for PCR were as follows:

Isg15: 5’- CTAGAGCTAGAGCCTGCAG -3’, 5’- AGTTAGTCACGGACACCAG -3’;

Mx1: 5’- GGTCCAAACTGCCTTCGTAA -3’, 5’- TTCAGCTTCCTTTTCTTGGTTT -3’;

Mx2: 5’- ACCAGGCTCCGAAAAGAGTT -3’, 5’- TCTCGTCCACGGTACTGCTT -3’;

Ifi204: 5’- GGGAGTGGAAAATGGCACAAC -3’, 5’- TCAGCACCATCACTTGTTTGG -3’;

Ifi27l2a: 5’- CTGTTTGGCTCTGCCATAGGA -3’, 5’- TTCCTGCACAGTGGACTTGAC -3’;

IL-6: 5’- CCGGAGAGGAGACTTCACAG -3’, 5’- TCCACGATTTCCCAGAGAAC -3’;

CCL2: 5’- AAGAGGATCACCAGCAGCAG -3’, 5’- TCTGGACCCATTCCTTCTTG -3’;

CCL17: 5’- AGTGGAGTGTTCCAGGGATG -3’, 5’- TGGCCTTCTTCACATGTTTG -3’;

CXCL10: 5’- CCAAGTGCTGCCGTCATTTTC -3’, 5’- GGCTCGCAGGGATGATTTCAA -3’;

CCL20: 5’- TTTTTGGGATGGAATTGGAC -3’, 5’- AGGTCTGTGCAGTGATGTGC -3’

CCL20: 5’- TTTTTGGGATGGAATTGGAC -3’, 5’- AGGTCTGTGCAGTGATGTGC -3’

Ly6g: 5’- GACTTCCTGCAACACAACTACC -3’, 5’- ACAGCATTACCAGTGATCTCAGT -3’

MPO: 5’- AGTTGTGCTGAGCTGTATGGA -3’, 5’- CGGCTGCTTGAAGTAAAACAGG -3’

ELANE: 5’- CAGGAACTTCGTCATGTCAGC -3’, 5’- AGCAGTTGTGATGGGTCAAAG -3’

CSF3R: 5’- CCTCACTTGAACTACACCCAGG -3’, 5’- CCCTTGGTACTGACAGTCGG -3’

C3: 5’- CCAGCTCCCCATTAGCTCTG -3’, 5’- GCACTTGCCTCTTTAGGAAGTC -3’

CXCL1: 5’- ACTGCACCCAAACCGAAGTC -3’, 5’- TGGGGACACCTTTTAGCATCTT -3’

CXCL2: 5’- CATCCAGAGCTTGAGTGTGACG -3’, 5’- GGCTTCAGGGTCAAGGCAAACT -3’

CXCL5: 5’- GTTCCATCTCGCCATTCATGC -3’, 5’- GCGGCTATGACTGAGGAAGG -3’

TNF: 5’- CCTGTAGCCCACGTCGTAG -3’, 5’- GGGAGTAGACAAGGTACAACCC -3’

Csf3: 5’- ATCCCGAAGGCTTCCCTGAGTG -3’, 5’- AGGAGACCTTGGTAGAGGCAGA -3’

### Primary culture of alveolar type II epithelial cells (AEC II)

Carefully excise the intact lung tissues of fetal mice (E19-20) and place them in pre-chilled ice-cold PBS. Remove residual tracheal and connective tissues to ensure only lung tissue is retained. Rinse the lungs twice with pre-chilled ice-cold PBS and cut into approximately 1 mm³ pieces using sharp ophthalmic scissors, adding 100 µL of antibiotics during the cutting process. Rinse the lung tissue pieces once with 0.25% trypsin (Gibco™; 25200072) (containing 0.01% DNase I (Sangon Biotech; A610099)) solution. Continue the digestion by pipetting the tissue with 0.25% trypsin (containing 0.01% DNase I) and incubate at 37°C in a shaking water bath for 10-20 minutes. Terminate the digestion process by adding an equal volume of DMEM/F12 medium (Gibco™; C11330500BT) containing 10% FBS (Gibco™; A5669701). Centrifuge the mixture at 1000 rpm for 8 minutes. Discard the supernatant and retain the cell pellet, which mainly consists of AECs and fibroblasts. Resuspend the cell pellet in 1 mL of 0.1% collagenase I solution (Sangon Biotech; A004214) and incubate at 37°C with 5% CO_2_ for 10-20 minutes (optimal duration 15 minutes). Terminate the digestion by adding an equal volume of DMEM/F12 medium containing 10% FBS and pipette thoroughly to mix. After filtering the cell suspension through a 200-mesh sieve, centrifuge at 1000 rpm for 5 minutes at low temperature, discard the supernatant, and collect the cell pellet, primarily containing AECs and minimal fibroblasts. Resuspend the cells in 3 mL of red blood cell lysis buffer and incubate at room temperature for 10 minutes to lysate the erythrocyte. Centrifuge at 1000 rpm for 5 minutes at low temperature, discard the supernatant and resuspend the cell pellet in DMEM/F12 medium containing 10% FBS. Incubate at 37°C with 5% CO_2_. After an hour, adherent cells are primarily fibroblasts, while non-adherent cells are mostly epithelial cells. Gently aspirate the culture medium, centrifuge at 4°C, 1000 rpm for 5 minutes and resuspend the pellet in fresh medium before seeding into new culture plates. Repeat this process one hour later and the remaining cells are primarily alveolar type II epithelial cells (AEC II). After 24 hours, change the culture medium and observe cell morphology and growth under an inverted microscope.

### Immunostaining and Confocal Microscopy

Cells were fixed with 4% FSB solution containing 4% paraformaldehyde (Servicebio; G1101), 4% sucrose (Sangon Biotech; A610498) in PBS for 30 min at room temperature (RT) followed by washing with PBS three times. Fixed cells were then exposed to 0.5 % Triton-X 100 and 5% serum in PBS for 30 minutes at RT for permeabilization and blocking. Next, cells were incubated with the primary antibody overnight at 4 ℃, diluted with 0.5% Triton X-100 and 5% serum in PBS. The primary antibodies used included the following: anti-TOM20 (1:500; Proteintech, # 11802-1-AP); anti-DNA (1:200; progen, # AC-30-10). Following three washes with PBS, the cells were incubated with Alexa 488 and 555 conjugated secondary antibodies (Invitrogen; 1:500, # A21422; #A11008) for 1 hour at RT, diluted with 0.5% Triton X-100 and 2% serum. The nuclei were then stained with DAPI (1:1000) for 10 min at RT. Finally, coverslips were mounted with anti-fade reagent. Images were acquired on a Zeiss LSM 980 confocal microscope with Airyscan technology. To maintain clarity and uniformity throughout the paper, some images have been pseudo-colored. As for lung tissue immunostaining, carefully excise the intact lung tissues of fetal mice (E19-20) and immerse them in pre-chilled ice-cold 4% paraformaldehyde (PFA). After 24 hours, the tissue was removed to 30% sucrose (Sangon Biotech; A610498), allowing the tissue to equilibrate until it sinks, which usually takes 48-72 hours. Take the precipitated lung tissue and blot dry the surface liquid with clean absorbent paper. Embed the tissue in OCT compound and rapidly freeze it using dry ice or liquid nitrogen. Cut 30 μm sections with a cryostat, then store the sections at −20 °C or proceed directly with subsequent immunofluorescence staining. Permeabilize the sections in 0.1% Triton X-100 for 10 min at RT followed by blocking with 3% BSA in PBS for an hour at RT. The antibody incubation process and the image acquisition were similar to that used for the cells.

### Detection of mtDNA content in cytosolic extracts

Freshly purified lung alveolar type II epithelial cells were resuspended in 500 μl buffer containing 150 mM NaCl, 50 mM HEPES pH 7.4 and 20 μg/ml digitonin (Sigma D141). The homogenates were incubated end-over-end for 10 min on the shaker to allow for selective plasma membrane permeabilization and then centrifuged three times at 800 g for 5 min to pellet intact cells. The supernatants were transferred to fresh tubes and centrifuged at 25,300 g for 10 min to pellet any remaining cellular chip, yielding cytosolic preparations free of nuclear, mitochondrial, and endoplasmic reticulum contamination. Samples were normalized for protein concentration using the BCA Protein Assay Kit (Epizyme Biomedical). DNA was extracted from equal amounts of cytosolic fractions using QIAquick Nucleotide Removal kit (QIAGEN) following the manufacturer’s instructions. Quantitative PCR was performed on pure cytosolic fractions using mtDNA primers: mouse mtND2 forward, 5′- ccatcaactcaatctcacttctatg-3′, and reverse, 5′- gaatcctgttagtggtggaagg-3′; mouse cytochrome b (Cytb) forward, 5′- cttcgctttccacttcatcttacc-3′, and reverse, 5′- ttgggttgtttgatcctgtttcg-3′; mouse Dloop1 forward, 5′- aatctaccatcctccgtgaaacc-3′, and reverse, 5′- tcagtttagctacccccaagtttaa-3′; For each independent sample, qPCR was performed in technical triplicates.

### Enzyme-Linked Immunosorbent Assay (ELISA)

Lungs were collected from fetal mice (E19-20) and placed in ice-cold PBS to remove any residual blood. The tissues were then weighed and homogenized in ice-cold PBS with protease inhibitor cocktail (Sigma-Aldrich, 11836170001) using a homogenizer, with the buffer volume approximately ten times the tissue weight (e.g.100 mg of tissue in 1 mL of buffer). The homogenate was centrifuged at 12,000 g for 10 minutes at 4 °C to pellet cellular debris. The protein concentration of the lysate was measured using the BCA Protein Assay Kit (Epizyme Biomedical). ELISA were performed according to the instructions of the mouse IL-1β ELISA kit (Proteintech; KE10003) and IFN-β ELISA kit (E-EL-M0033) and the final concentrations were normalized to respective protein concentration.

### Blood cell smear

Blood samples were collected from mice via using the decapitation method using heparinized capillary tubes to prevent coagulation. The original sentence is grammatically correct, but it can be slightly improved for clarity and conciseness. A small drop of blood (5-10 µL) was then placed near one end of a clean microscope slide, and a thin, even smear was created by quickly and smoothly pulling a spreader slide across the stationary slide at a 30-45 degree angle. The blood smears were air-dried completely at room temperature, fixed in methanol for 2-3 minutes, and stained using Wright-Giemsa stain solution by immersing the slides for 10 minutes, rinsing with distilled water, and air drying. The stained blood smears were examined under a light microscope at 100x magnification using oil immersion. Differential white blood cell counts were performed by identifying and counting 100 leukocytes per slide, including neutrophils, lymphocytes and monocytes, and the total number of each cell type was recorded.

### Quantification and statistical analysis

Student’s t-test was used to analyze differences between two groups, and One-way or Two-way ANOVA was used to analyze intergroup differences. P-values less than 0.05 were considered statistically significant. The analysis was performed using GraphPad Prism 8 (GraphPad software). Densitometry results of western blotting were quantified using ImageJ software. All data were presented as mean ± SEM and other details such as the number of replicates and the level of significance was mentioned in figure legends and supplementary tables.

## Acknowledgments

This work is supported by the National Natural Science Foundation of China: JSK (32071137, 92054103). Funding for Scientific Research and Innovation Team of The First Affiliated Hospital of Zhengzhou University: JSK (ZYCXTD2023014).

## Author contributions

DHL designed the experiments, performed experiments, analyzed the data and wrote manuscript. FL did preliminary exploration and part of the mice experiments. RZY discussed the project and provided help in the experiments design and data analysis. ZBW performed zebrafish experiments and data analysis. XYM and SML provided help in the experiments and manuscript writing. WXL, JKL, DDW and RYW provided help in the experiment conduction. SAL and PPL provided help in the experiments design. JSK developed the idea, directed the project and review the paper. All authors participated in discussions.

---

**Figure S1:**
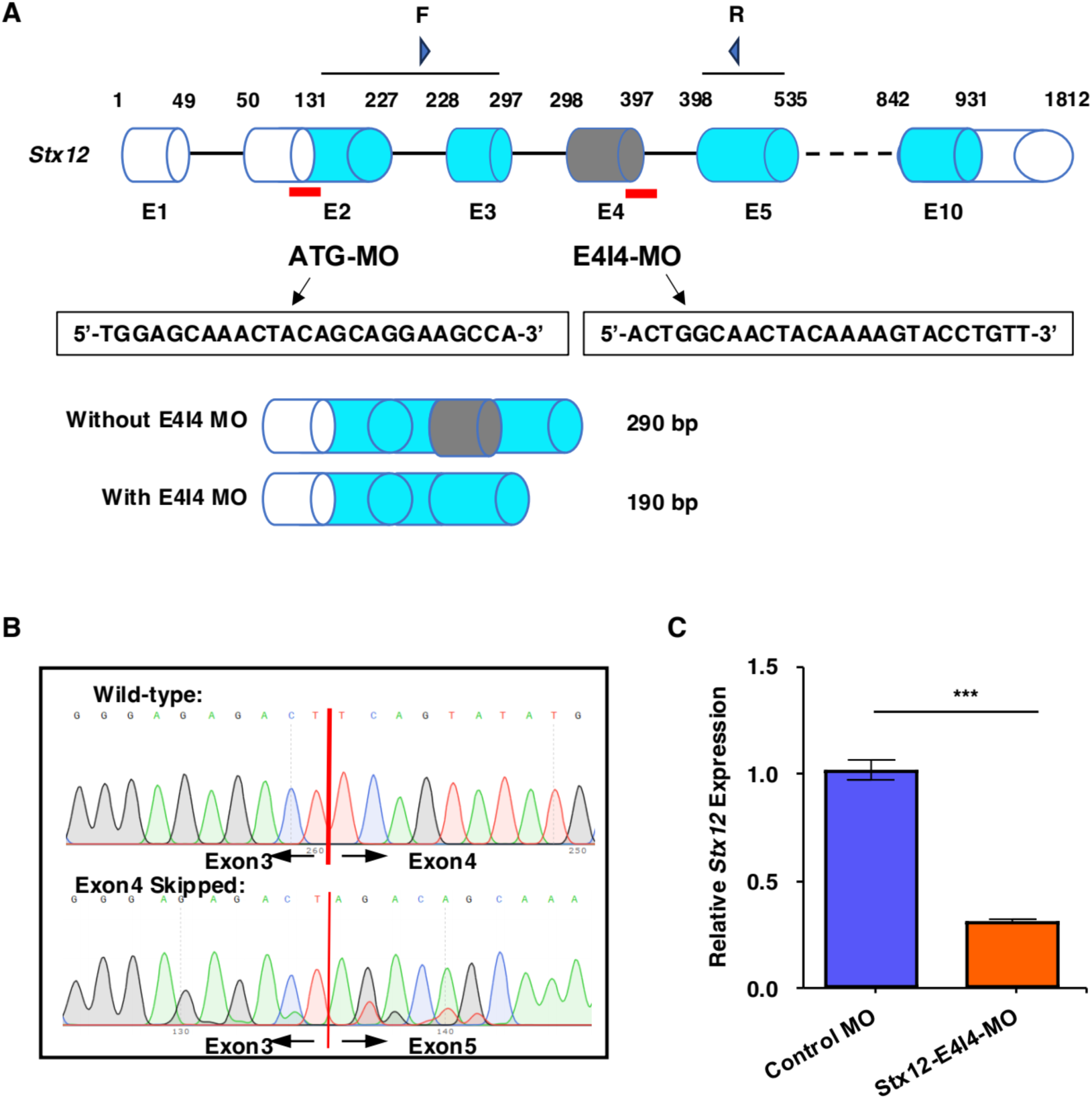
Design and effectiveness of Zebrafish *Stx12* Morpholino. **A.** Schematic of *Stx12* antisense morpholino oligonucleotide (MO) targeting ATG and the exon-intron junction (E4). Primers F and R interrogate the presence of wild type (non-mutant) transcripts or those in which exon 4 has been skipped. **B.** Sanger sequencing of both the wild type band and the exon 4-skipped band validating the wild type sequence and the exon 4-skipped sequence. **C.** Quantitative measurements of *stx12* expression levels measured by qRT-PCR. MO-targeted down-regulation of *stx12*.

**Figure S2:**
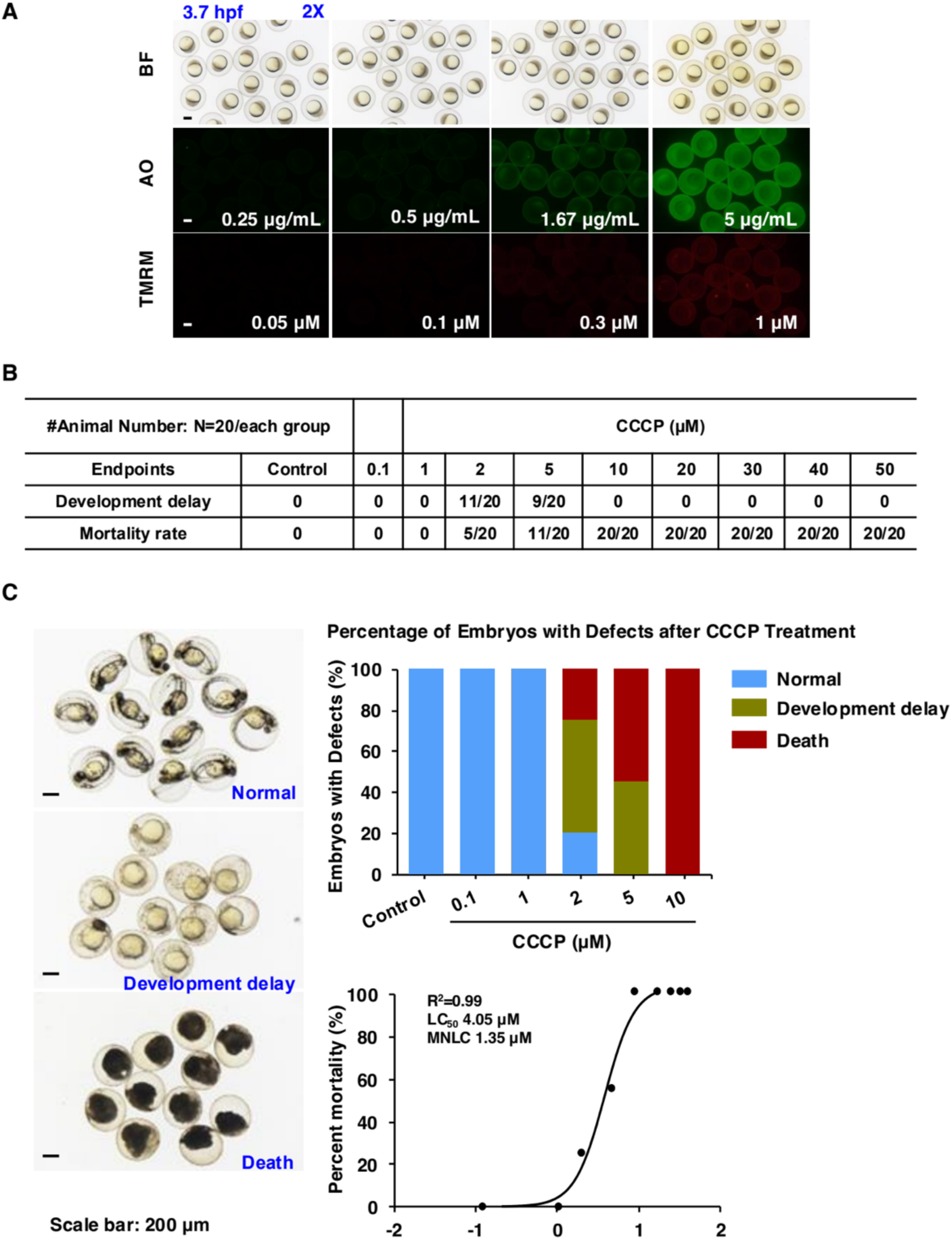
TMRM Staining & AO Staining: dose-dependent study and the investigation of maximum non-lethal concentration (MNLC) and LC50 determination of CCCP. **A.** Representative images of zebrafish embryos stained with different concentrations of TMRM (0.05 μM, 0.1 μM, 0.3 μM, 1 μM) and AO (0.25 μg/mL, 0.5 μg/mL, 1.67 μg/mL and 5 μg/mL). TMRM (Tetramethylrhodamine methyl ester) indicates mitochondrial membrane potential and AO (Acridine Orange) indicates apoptosis. Scale bar: 300 μm. **B.** Statistic number of zebrafish situations observed after CCCP treatment. **C.** Representative images under different conditions and statistical analysis of the data from A. Scale bar: 200 μm.

**Figure S3.**
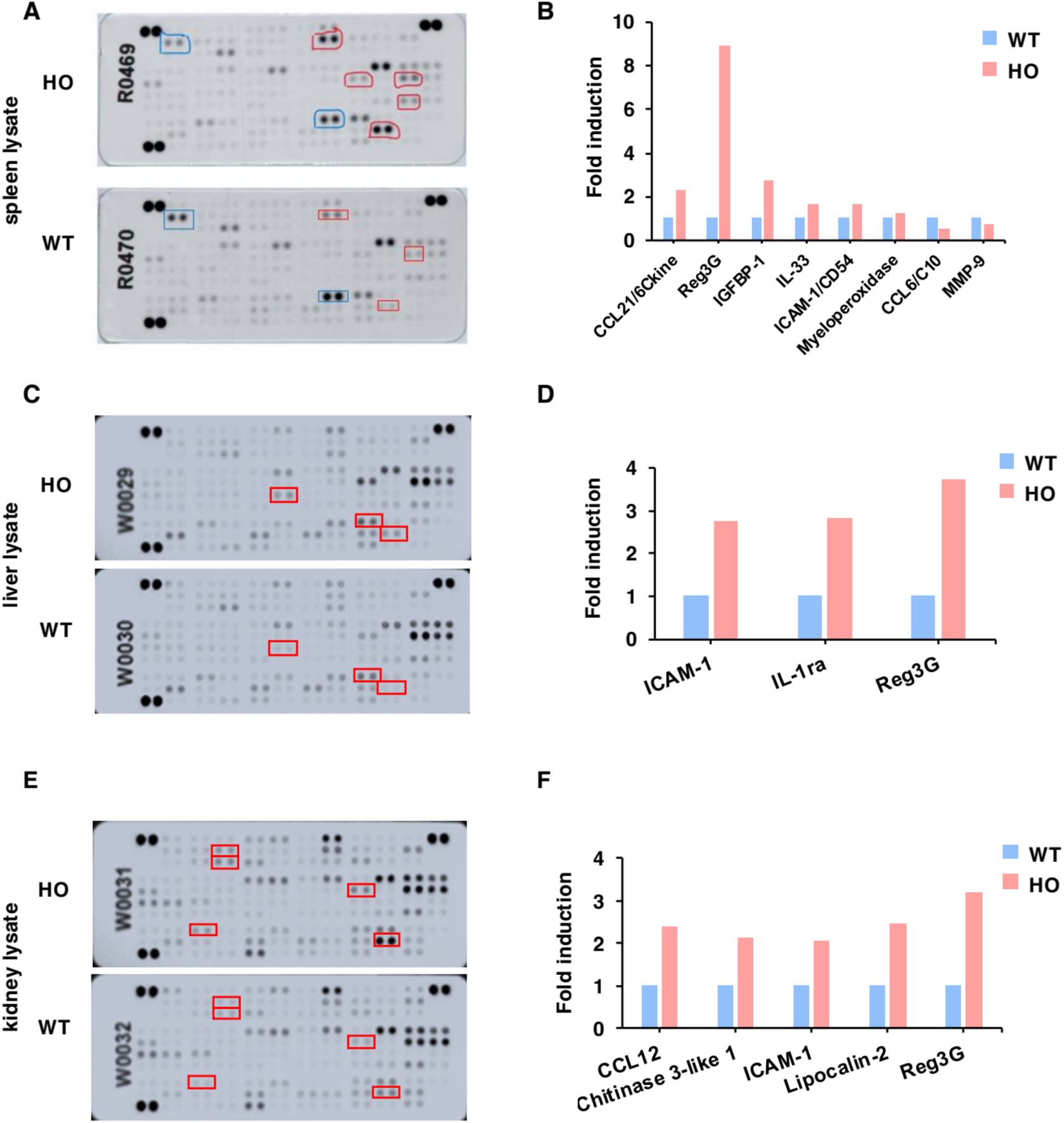
The cytokine level of brain lysate, spleen lysate, liver lysate and kidney lysate. **A, B.** The cytokine kit displays of spleen lysate from *Stx12*^−/−^ mice (HO) and the littermate controls (WT) (R&D Systems, Catalog#410-MT) and quantitative analysis. **C, D.** The cytokine kit displays of liver lysate from *Stx12*^−/−^ mice (HO) and the littermate controls (WT) (R&D Systems, Catalog #410-MT) and quantitative analysis. **E, F.** The cytokine kit displays of kidney lysate from *Stx12*^−/−^ mice (HO) and the littermate controls (WT) (R&D Systems, Catalog #410-MT) and quantitative analysis.

**Figure S4.**
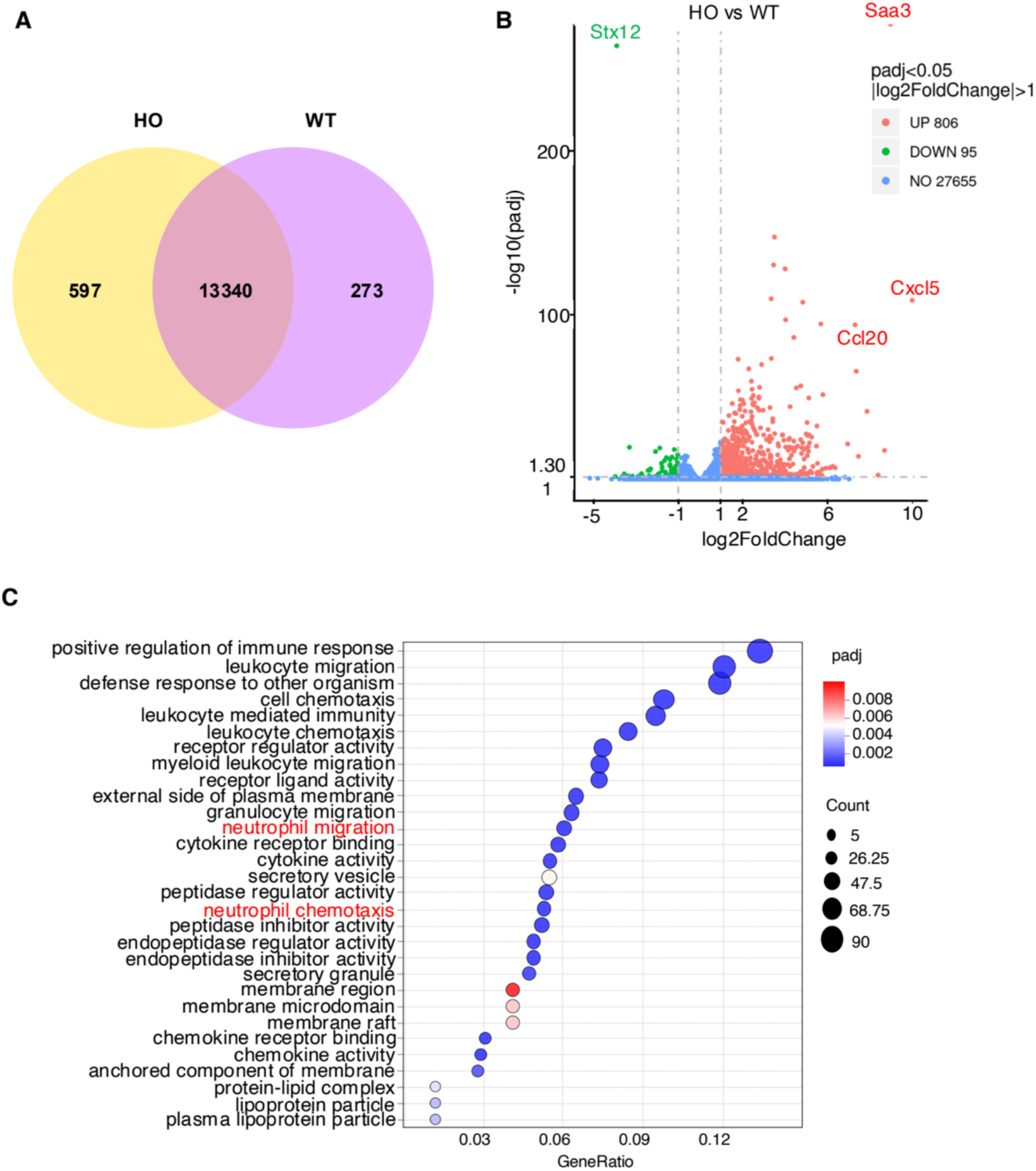
Differential express between the lung tissues from *Stx12^−/−^* mice (HO) versus the littermate controls (WT, *Stx12*^+/+^). **A.** The number of co-expression gene between the lung tissues from *Stx12^−/−^* mice (HO) versus the littermate controls (WT, *Stx12*^+/+^). **B.** Volcano plot of differentially expressed genes in Stx12^−/−^ mice (HO) versus the littermate controls (WT, *Stx12*^+/+^) lung tissue. **C.** GO enrichment bubble plot of differentially upregulated genes in Stx12^−/−^ mice lung tissue. The bubble plot illustrates the significantly enriched Gene Ontology (GO) terms for biological processes (BP), molecular function (MF) and cellular component (CC) related to the differentially upregulated genes. The x-axis represents the Gene Ratio while the y-axis lists the corresponding GO terms. The size of each bubble reflects the number of upregulated genes associated with each GO term, and the color of the bubbles denotes the statistical significance (adjusted p-value) of the enrichment. All results are presented as mean ± SEM; statistical significance was assessed by Student’s t-test.

**Figure S5.**
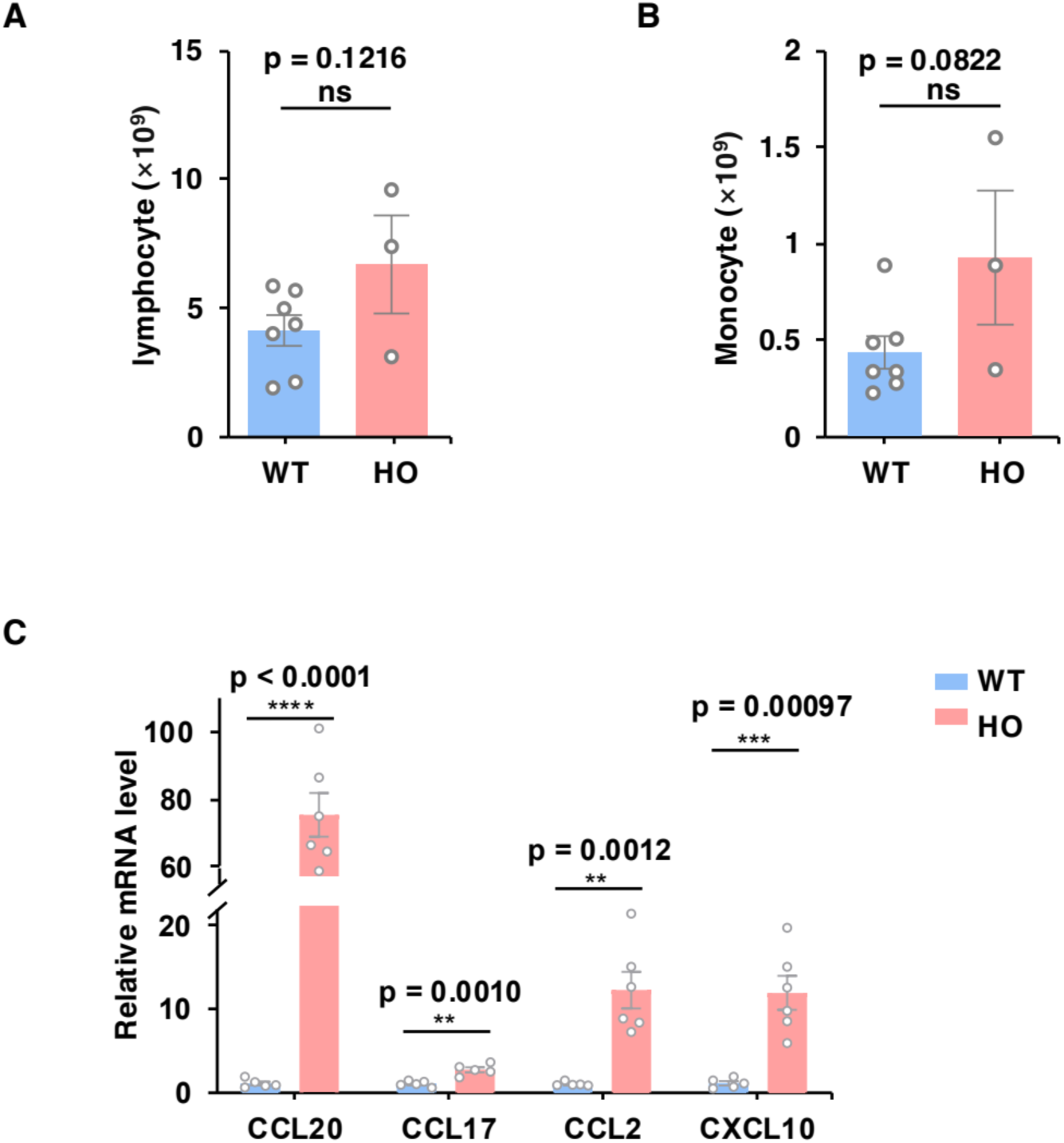
The number of lymphocyte and monocyte in the blood from Stx12^−/−^ mice (HO) and the littermate controls (WT, *Stx12*^+/+^) and the relative mRNA level of chemokines. **A.** Quantification of lymphocytes in the blood from the littermate controls (WT, *Stx12*^+/+^) and *Stx12^−/−^* mice. **B.** Quantification of monocytes in the blood from the littermate controls (WT, *Stx12*^+/+^) and *Stx12^−/−^* mice. **C.** The relative mRNA expression levels of chemokines CCL20, CCL17, CCL2 and CXCL10 by qRT-PCR from lung tissues of *Stx12^−/−^* (HO, n = 6) and the littermate controls (WT, *Stx12*^+/+^) (n = 5) mice. The mRNA expressions of target genes were normalized to β-actin. Results are presented as mean ± SEM; statistical significance was assessed by Student’s *t*-test.

